# Psychedelic compounds directly excite 5-HT_2A_ Layer 5 Pyramidal Neurons in the Prefrontal Cortex through a 5-HT_2A_ Gq -mediated activation mechanism

**DOI:** 10.1101/2022.11.15.516655

**Authors:** Gavin P. Schmitz, Yi-Ting Chiu, Gabriele M. König, Evi Kostenis, Bryan L. Roth, Melissa A. Herman

## Abstract

Psilocin, the active compound in *Psilocybe sp*. mushrooms, is a serotonergic psychedelic that has recently gained renewed interest due to its potential as a therapeutic tool. Despite promising clinical findings, the underlying signaling mechanisms and brain region-specific effects of psilocin and other psychedelic drugs remain unclear. Psilocin, like other psychedelic compounds, is an agonist at many serotonin and other biogenic amine receptors; however, activation of serotonin (5-Hydroxytryptamine, or 5-HT) 2A receptors (5-HT_2A_Rs) is understood as the main molecular target for the psychoactive effects in animals and humans. 5-HT_2A_Rs are abundantly expressed in the prefrontal cortex (PFC); however, the biochemical actions of psilocin on PFC neurons remain poorly understood. In this study, we used *in vitro* slice electrophysiology to examine how psilocin acutely alters the activity and electrophysiological properties of layer 5 pyramidal neurons in the mouse PFC. Focal application of psilocin (10*μ*M) onto nonspecified Layer 5 Pyramidal neurons in the prelimbic PFC of C57BL/6J mice produced variable effects on firing (increase, decrease, or no change). 5-HT_2A_R layer 5 pyramidal neurons in the mouse prelimbic PFC were identified via labeling in a 5-HT2A-ERT2-Cre mouse crossed with an Ai9 tdTomato reporter. Focal application of psilocin increased firing in all identified 5-HT_2A_R neurons but did not result in any significant changes in synaptic transmission. Overall, the results demonstrate that psilocin evokes strong firing changes in the PFC that are 5-HT_2A_R and G*α*_q_ dependent, thereby providing valuable insights into the effects of psilocin on a brain region implicated in mediating psychedelic drug actions.

## 1. Introduction

Psychedelic drugs are compounds that induce changes in cognition, emotion, and perception via activation of 5-HT_2A_ serotonin (5-hydroxytryptamine; 5-HT) receptors^1, 2^. Recently, there has been renewed interest in psychedelic compounds as potential therapeutics for many neuropsychiatric disorders^3^. Psilocybin, the prodrug to the active compound psilocin found in *Psilocybe sp*. mushrooms, has recently been granted breakthrough drug status by the Food and Drug Administration due to its potential as a treatment for treatment-resistant depression and anxiety^4, 5, 6^. The effects of psilocybin in these Phase II clinical trials are rapid, robust and persistent^4, 5, 7^.

Psychedelics are generally classified by chemical structure (e.g. tryptamines; ergolines and phenylisopropylamines), and each psychedelic drug has a robust pharmacology with activities at many serotonin and other biogenic amine receptors^8-12^. 5-HT_2A_ receptor agonism has been shown to be crucial in mediating the hallucinogenic effects of psychedelics in both animal^13-15^ and human studies^16-19^. 5-HT_2A_ receptors are members of the G protein-coupled receptors (GPCR) superfamily and canonically signal through the G*α*_q_ family of heterotrimeric G proteins activating phospholipase Cb and many other down-stream effector systems ^20, 21, 2^. 5-HT_2A_ receptors also recruit β-arrestin (βArr) proteins *in vitro*^22^ and the 5-HT_2A_R is apparently complexed with βArr1 and βArr2 in cortical neurons *in vivo*^*23*^. Recently, there has been considerable interest in creating ligands with biased signaling profiles with the aim of creating non-hallucinogenic 5-HT_2A_R agonists for potential therapeutic application^24-27^. However, it remains unknown whether the effects of existing compounds under investigation are due to G*α*_q_ signaling, β-arrestin recruitment, or alternative signaling pathways^28-30^.

Additionally, the brain region-specific effects of psychedelic drugs like psilocin remain poorly understood despite their therapeutic promise, particularly in pre-clinical animal models. Positron emission tomography studies in humans demonstrate that the psychedelic effects of psilocin correlate with PFC 5-HT_2A_R occupancy^31^. In clinical trials on patients with treatment resistant depression and nicotine addiction, patients reported that psilocin’s ability to modulate a person’s self-focus, reduce negative emotional processing, and strengthen social functioning^32, 33^ were crucial to psilocin’s therapeutic effect. The PFC plays an important role in both the control of high-level cognitive and emotional processes and enabling a sense of self^34^. Additionally, maladaptation of PFC functioning has been linked to depression^35^. In light of the above, understanding the effect of psilocin on PFC neurons is essential to understanding psilocin’s many actions.

Here, we comprehensively examined the acute effects of psilocin and the selective 5-HT_2A_R agonist 25-CN-NBOH on layer 5 pyramidal neurons in the Prelimbic region of the Prefrontal Cortex. We observed that both drugs significantly increase firing in layer 5 pyramidal neurons that express the 5-HT_2A_R. Application of the 5-HT_2C_ antagonist RS102221 (5mM) prior to psilocin administration failed to block the increased firing with application of psilocin, while application of the 5-HT_2A_ antagonist M100907 (100nM) prior to psilocin administration did prevent increased firing. Furthermore, application of the G*α*_q_ inhibitor FR100907 (1*μ*M) prior to psilocin administration also blocked the increase in firing with psilocin administration. These results demonstrate that psilocin selectively increases the activity of 5-HT_2A_ Layer 5 PFC pyramidal neurons via a 5-HT_2A_ receptor-dependent mechanism. Additionally, this effect is dependent upon signaling via the G*α*_q_ G protein.

## 2. Materials and Methods

### 2.1 Animal Use and Treatment

All experiments were conducted in male and female adult mice (19-30g). Experiments were performed in either C57BL6/J mice or transgenic 5-HT_2A_R-eGFP-CreERT2 mice crossed with an Ai9 reporter mouse line. This 5-HT_2A_R-eGFP-CreERT2 mouse expresses 5-HT_2A_R tagged with GFP as described^36^ (Chiu et al, *in preparation*). To induce recombination, tamoxifen (100 mg/kg, i.p) was administered by injection once daily over 4 days in mice p35 or older, when mice reach sexual maturity and receptor expression patterns reflect that of the adult mouse. Mice were monitored for at least 1-2 weeks after tamoxifen injections before use. Mice were bred at the University of North Carolina at Chapel Hill and maintained with ad libitum access to food, water, and environmental enrichment (nesting materials). All procedures were carried out in accordance with the NIH Guide for the Care and Use of Laboratory Animals and as approved by the Institutional Animal Care and Use Committee.

### 2.2 Immunohistochemistry

2A-eGFP-Cre^ERT2^ mice were crossed with the Cre reporter Ai9 strain (#007909; Jackson Labs, Bar Harbor, ME). Mice were Intraperitoneally injected with 100mg/kg tamoxifen (TAM) from p39 to p42 and incubated for another 1-2 weeks and then brains were dissected out at p56. Brain sections through frontal lobe region were stained with anti-goat-GFP antibody (1:1000, Rockland, #600-101-215) followed by donkey anti-rabbit-Alexa Fluor® 488 (1:1000, Jackson Immunoresearch, West Grove, PA). Brain section images were taken by Olympus VS120 virtual slide microscope (Olympus, Tokyo, Japan) to see 5-HT_2A_R-GFP fusion protein and tdTomato signal under 10X magnification.

### 2.3 Electrophysiological Recording

Male and female WT and transgenic 5-HT_2A_R-eGFP-CreERT2xAi9 mice were rapidly decapitated, and brains placed in a beaker containing ice-cold high sucrose solution (in mM): sucrose 206.0; KCl 2.5; CaCl_2_ 0.5; MgCl_2_ 7.0; NaH_2_PO_4_ 1.2; NaHCO_3_ 26; glucose 5.0; HEPES 5. Coronal sections (300*μ*m) were sliced on a vibrating microtome (Leica VT1000S, Leica Microsystems, Buffalo Grove, IL) and incubated in an oxygenated (95% O_2_/5% CO_2_) artificial cerebrospinal fluid (aCSF) composed of (in mM): NaCl 130; KCl 3.5; Glucose 10; NaHCO3 24; MgSO4-7H2O 1.5; NaH2PO4-H2O 1.26; CaCl 2.0 for 30 min at 37°C, followed by 30min at room temperature (RT, 21-22°C) for equilibration. Patch pipettes (4-6 MΩ; King Precision Glass Inc., Claremont, CA) were filled with internal solution (mM: potassium gluconate 145; EGTA 5; MgCl2 5; HEPES 10; Na-ATP 2; Na-GTP 0.2 (for EPSC recordings) or potassium chloride [KCl] 145; EGTA 5; MgCl_2_ 5; HEPES 10; Na-ATP 2; Na-GTP 0.2 (for IPSC recordings)). Data were acquired with a Multiclamp 700B amplifier (Molecular Devices, Sunnyvale, CA), low-pass filtered at 2-5 kHz, digitized (Digidata 1550B; Molecular Devices) and stored using pClamp 10 software (Molecular Devices). Series resistance was continuously monitored with a hyperpolarizing 10mV pulse; cells with axis resistance >20 MΩ were excluded from the data set. 5HT_2A_R expressing neurons were identified via Tdtomato expression using fluorescent optics and brief (<2s) episodic illumination as previously described^37^. During voltage clamp recording (V_hold_ = -70mV), electrophysiological properties of cells were determined by pClamp 10 Clampex software using a 10mV pulse delivered after breaking into the cell. Current Clamp recordings (I_hold_ = 70pA) were performed immediately after breaking into cell to determine the firing type of each neuron included in the data sets. To isolate currents mediated by glutamate receptors, recordings were conducted in the presence of the GABA receptor antagonists SR-95531 (gabazine, GBZ, 20 μM) and CGP 52432 (CGP, 1 μM). Isolation of GABA_A_ sIPSCs was achieved through selective pharmacological blockade with the glutamate receptor blockers 6,7-dinitroquinoxaline-2,3-dione (DNQX, 20*μ*M) and DL-2-amino-5-phosphonovalerate (AP-5, 50*μ*M) and the GABA_B_ receptor antagonist CGP55845 (1*μ*M). The sodium channel blocker tetrodotoxin citrate (TTX, 1 μM) was added to the bath for mEPSC recordings.

### 2.4 Cell culture

HEK293T cells were obtained from ATCC (Manassas, VA). Cells were maintained, passaged, and transfected in Dulbecco’s Modified Eagle’s Medium containing 10% fetal bovine serum (FBS), 100 Units/mL penicillin, and 100μg/mL streptomycin (Gibco-ThermoFisher, Waltham, MA) in a humidified atmosphere at 37°C and 5% CO2. After transfection, cells were plated in DMEM containing 1% dialyzed FBS, 100 Units/mL penicillin, and 100μg/mL streptomycin for BRET assays.

### 2.5 BRET2 assays

Cells were plated either in six-well dishes at a density of 700,000–800,000 cells/well, or 10-cm dishes at 7–8 million cells/dish. Cells were transfected 2–4 hours later, using a 1:1:1:1 DNA ratio of receptor:Gα-RLuc8:Gβ:Gγ-GFP2 (100 ng/construct for six-well dishes, 1000 ng/construct for 10-cm dishes). Transit 2020 (Mirus Biosciences, Madison, WI) was used to complex the DNA at a ratio of 3 μL Transit/μg DNA, in OptiMEM (Gibco-ThermoFisher, Waltham, MA) at a concentration of 10 ng DNA/μL OptiMEM. The next day, cells were harvested from the plate and plated in poly-D-lysine-coated white, clear bottom 96-well assay plates (Greiner Bio-One, Monroe, NC) at a density of 30,000–50,000 cells/well.

One day after plating in 96-well assay plates, white backings (Perkin Elmer, Waltham, MA) were applied to the plate bottoms, and growth medium was carefully aspirated and replaced immediately with 50 μL of assay buffer (1x HBSS + 20 mM HEPES, pH 7.4) containing freshly prepared 50 μM coelenterazine 400a (Nanolight Technologies, Pinetop, AZ). After a five-minute equilibration period, cells were treated with 50 μL of drug (2X) for an additional 5 minutes. Plates were then read in a Pherastar FSX microplate reader (BMG Labtech Inc., Cary, NC) with 395 nm (RLuc8-coelenterazine 400a) and 510 nm (GFP2) emission filters, at 1 second/well integration times. BRET2 ratios were computed as the ratio of the GFP2 emission to RLuc8 emission. Results are from at least three independent experiments, each performed in duplicate. Data were normalized to 5-HT stimulation and analyzed using nonlinear regression “log(agonist) vs. response” in Prism 9 (Graphpad Software Inc., San Diego, CA).

### 2.6 Drugs

Psilocin was purchased from Cayman chemical (Ann Arbor, MI). RS102221 was purchased from Tocris Bioscience (Ellisville, MO). M100907 was purchased from Sigma-Aldrich. FR900359 was obtained from the laboratory of Evi Kostenis using the extraction process described previously^38^. Stock solutions were prepared in ultra-pure water or DMSO, stored at 20°C and diluted to final experimental concentration in aCSF on the day of testing. Doses for electrophysiology experiments were chosen based on their ex vivo effects in PFC neurons from prior reports.

### 2.7 Experimental design and statistical analysis

All data analyses and visualizations were completed with Prism 9 (GraphPad, La Jolla, CA). Membrane characteristics were represented as mean ± SEM. Frequency and amplitude of spontaneous inhibitory postsynaptic currents (sIPSCs) and spontaneous excitatory postsynaptic currents (sEPSCs) were analyzed and manually confirmed using a semiautomated threshold-based detection software (Mini Analysis, Synaptosoft Inc.). sIPSC and sEPSC characteristics were determined from baseline and experimental drug application containing a minimum of either 60 events or spanning a duration of at least 3 minutes. Event data were represented as mean ± SEM and analyzed for normality. Data were analyzed for independent significance using a paired t-test if normally distributed or Wilcoxon test if not normally distributed. All concentration-response curves were fit to a three-parameter logistic equation in Prism 9 (Graphpad, La Jolla, CA). BRET2 concentration-response curves were analyzed by normalizing to a reference agonist for each experiment. Efficacy (Emax) calculations were performed according to Kenakin and colleagues^39^: stimulus-response amplitudes (net BRET2) were normalized to the maximal responding reference agonist (maximal system response). EC50 and Emax values were estimated from the simultaneous fitting of all biological replicates.

## 3. Results

### 3.1 Psilocin has differential effects on Layer 5 Pyramidal neurons in the Prefrontal Cortex of C57BL/6J mice

To investigate how acute psilocin exposure alters the firing of layer 5 pyramidal neurons, we performed whole-cell current-clamp recordings in the Prelimbic (PrL) PFC to measure changes in firing. Layer 5 pyramidal neurons were identified based on morphology and membrane characteristics (**Figure 1A**). Focal application of psilocin (10*μ*M) had variable effects on layer 5 pyramidal neurons in the PrL PFC. In most neurons, psilocin (10*μ*M) produced a significant increase in firing frequency, defined as a change greater than 120% of baseline (n=10 cells. Paired t-test *p<0.05. **Figure 1B, 1F**). In ∼30% of layer 5 pyramidal neurons in the PrL PFC, psilocin produced a decrease in firing frequency, defined as a change less than 80% of baseline (**Figure 1C, 1G**). In the remaining portion of layer 5 pyramidal neurons in the PrL PFC, psilocin produced no change in firing frequency, defined as a change between 80 – 120% (**Figure 1D, 1H**). Comparison of the membrane characteristics of the cells in these three groups did not show any significant differences in membrane capacitance (Cm), membrane resistance (Rm), or Tau (**Table 1**).

**Table 1:** Membrane characteristics of layer 5 pyramidal neurons shown as value +/-SEM.

|  | Cm | Rm | Tau | Hold (pA) | Vm (mv) |
| --- | --- | --- | --- | --- | --- |
| Increase Group | 112.5+/-12.37 | 177.7+/-65.79 | 2.01+/-1.101 | -49.90+/-38.18 | -63.86+/-2.473 |
| Decrease Group | 111.5+/-13.86 | 136.9+/-21.11 | 1.981+/-0.4542 | -237.9+/-212.5 | -61.10+/-1.219 |
| No Change Group | 120.7+/-10.8 | 108.5+/-8.256 | 2.55+/-0.45 | -46.25+/-32.75 | -64.68+/-2.393 |

**Figure 1:**
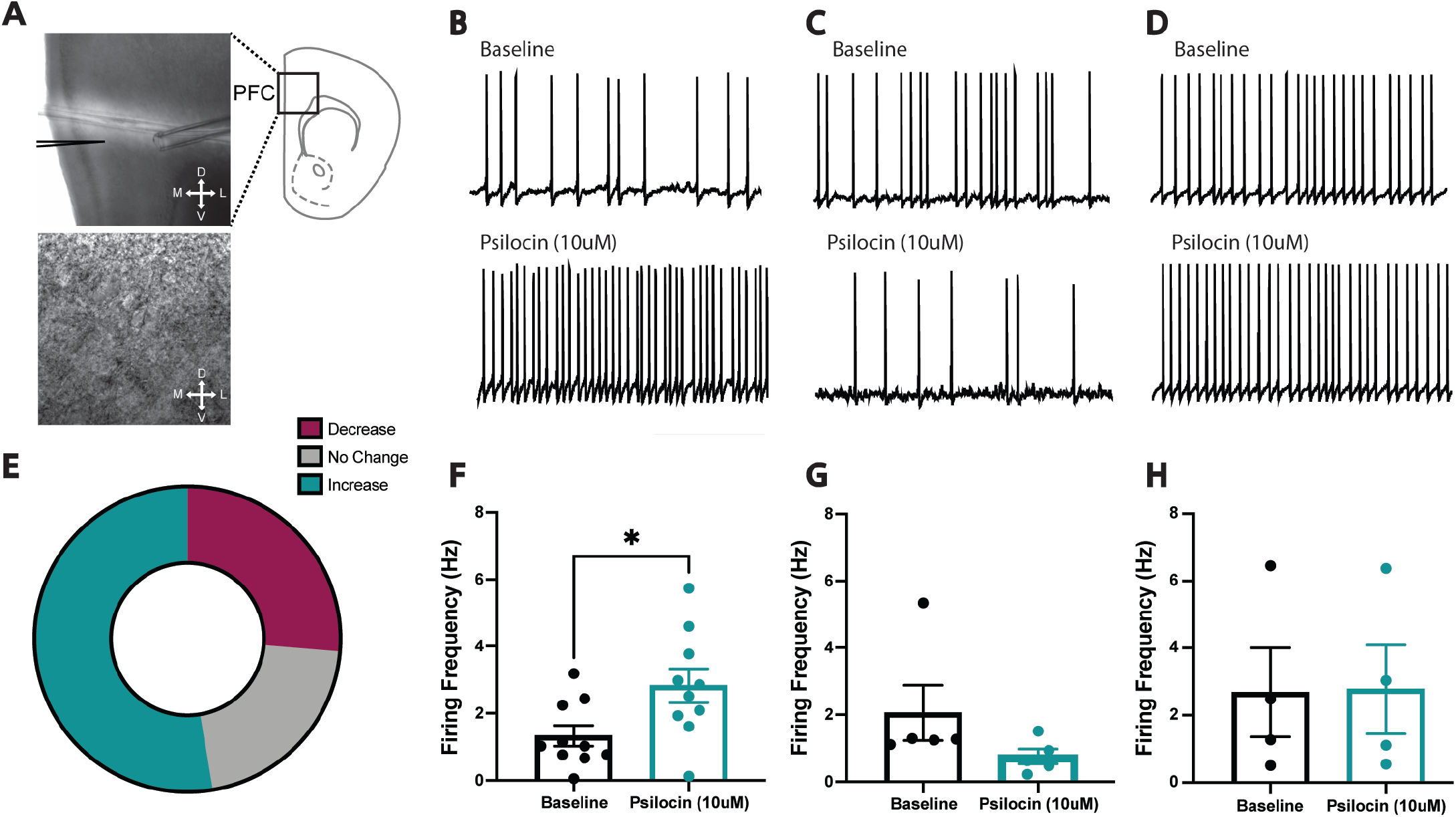
**(a)** Representative L5P neuron in PFC from C57BL/6J mouse and its location within slice. Representative recording of L5P neuronal firing after psilocin (10*μ*M) showing significant increase in frequency **(b)**, decrease **(c)**, and no change **(d). (e)** Graphical depiction of differential neuronal responses to psilocin (10*μ*M). **(f)** Averaged psilocin effects on firing in L5P neurons in the increase group (Data represented as mean ± SEM, n=10. Paired t-test *p<0.05). **(g)** Averaged psilocin effects on firing in L5P neurons in the decrease group (Data represented as mean ± SEM, n=5). (h) Averaged psilocin effects on firing in L5P neurons in the no change group. (Data represented as mean ± SEM, n=4).

### 3.2 5HT_2A_R-tdTomato-Reporter Mouse allows visualization of 5-HT_2A_R-expressing cells

To validate 5-HT_2A_R expression throughout the prefrontal cortex, we crossed our 5-HT_2A_R-eGFP-CreERT2 mouse line with an Ai9 tdTomato-reporter mouse line. This mouse line is useful for cre-dependent expression in selected 5-HT_2A_R-circuits and provides a representative map of 5-HT_2A_R-expressing cells, 5-HT_2A_R axon projections, and terminal fields (Chiu et al, *in preparation*). We characterized 5-HT_2A_R expression across the full brain of these mice and found robust 5-HT_2A_R-induced tdTomato expression through the prelimbic region of the prefrontal cortex (**Figure 2**) (Chiu et al, *in preparation*).

**Figure 2:**
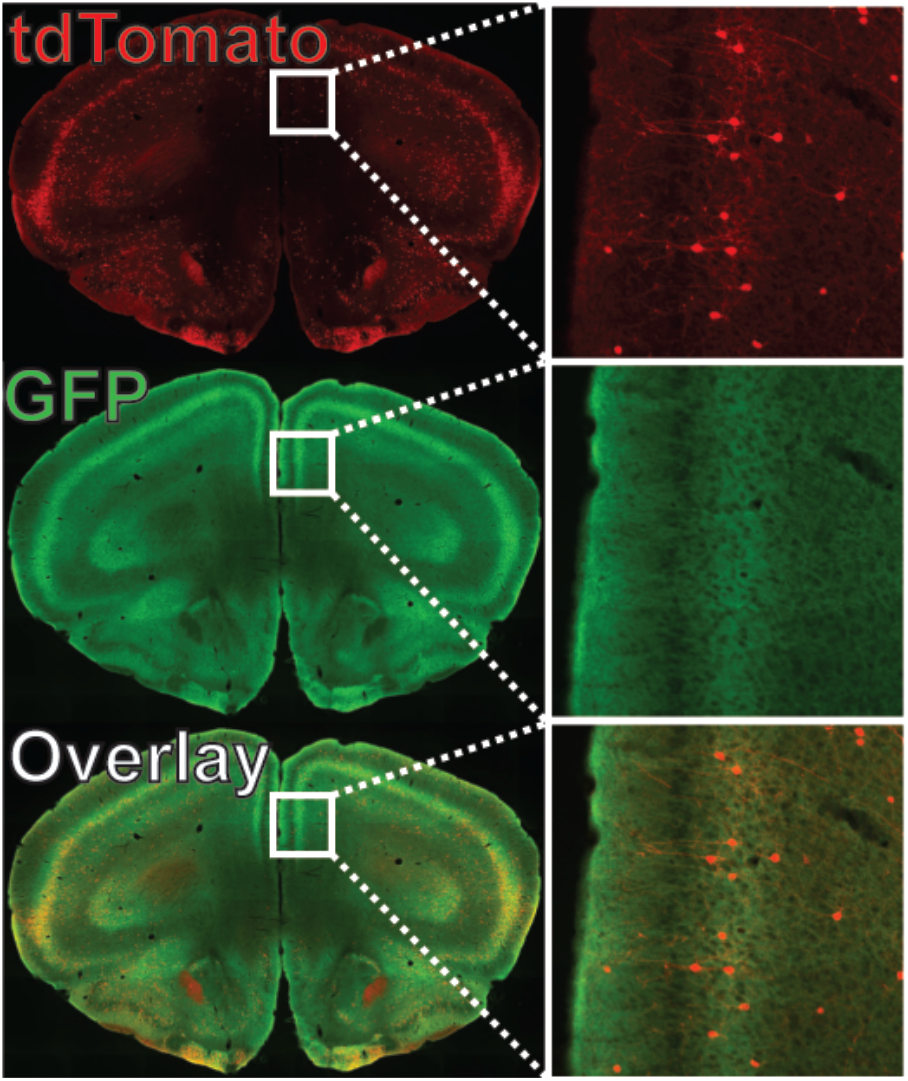
Visualization of tamoxifen induced expression (red) and 5-HT_2A_R (green) in the PFC of 5-HT_2A_R-eGFP-CreERT2xAi9 model.

### 3.3 Psilocin and NBOH-2C-CN increase firing in 5HT_2A_R layer 5 pyramidal neurons in the Prefrontal Cortex without impacting excitatory or inhibitory synaptic transmission

To specifically examine the effects of psilocin on 5-HT_2A_R expressing layer 5 pyramidal neurons in the PFC, we performed whole-cell current-clamp recordings in the PrL PFC to measure changes in firing in tdTomato-expressing neurons. Here, 5-HT_2A_R layer 5 pyramidal neurons were identified based on morphology and tdTomato expression (**Figure 3A**). Focal application of psilocin (10*μ*M) significantly increased firing frequency in all 5-HT_2A_R expressing neurons (n=13 cells. Paired t-test *p<0.05; **Figure 3B,C**). When firing frequencies were normalized to baseline values, firing frequency increased to ∼200% (n=13 cells. Wilcoxon signed rank test ***p (two tailed) <0.001; **Figure 3D**). To measure changes in excitatory glutamatergic synaptic transmission, we performed whole-cell voltage-clamp recordings in 5-HT_2A_R pyramidal neurons in the PrL PFC. Focal application of psilocin (10*μ*M) produced no changes in spontaneous excitatory post-synaptic current (sEPSC) frequency either in raw values (n=12 cells. **Figure 3E,F**) or normalized values (n=12 cells. **Figure 3G**). Similarly, psilocin produced no changes sEPSC amplitude either in raw values (**Figure 3H**) or normalized values (**Figure 3I**). To measure changes in glutamate release isolated to the pre-synaptic terminals, we performed recordings of miniature excitatory post-synaptic currents (mEPSCs). Application of psilocin (10*μ*M) produced no changes in mEPSC frequency or amplitude (**Figure S1A-E**). To measure changes in inhibitory synaptic transmission, we performed whole-cell voltage-clamp recordings in 5-HT_2A_R pyramidal neurons in the PrL PFC. Focal application of psilocin (10*μ*M) produced no changes in spontaneous inhibitory post-synaptic current (sIPSC) frequency or amplitude (**Figure S2A-E**).

**Figure 3:**
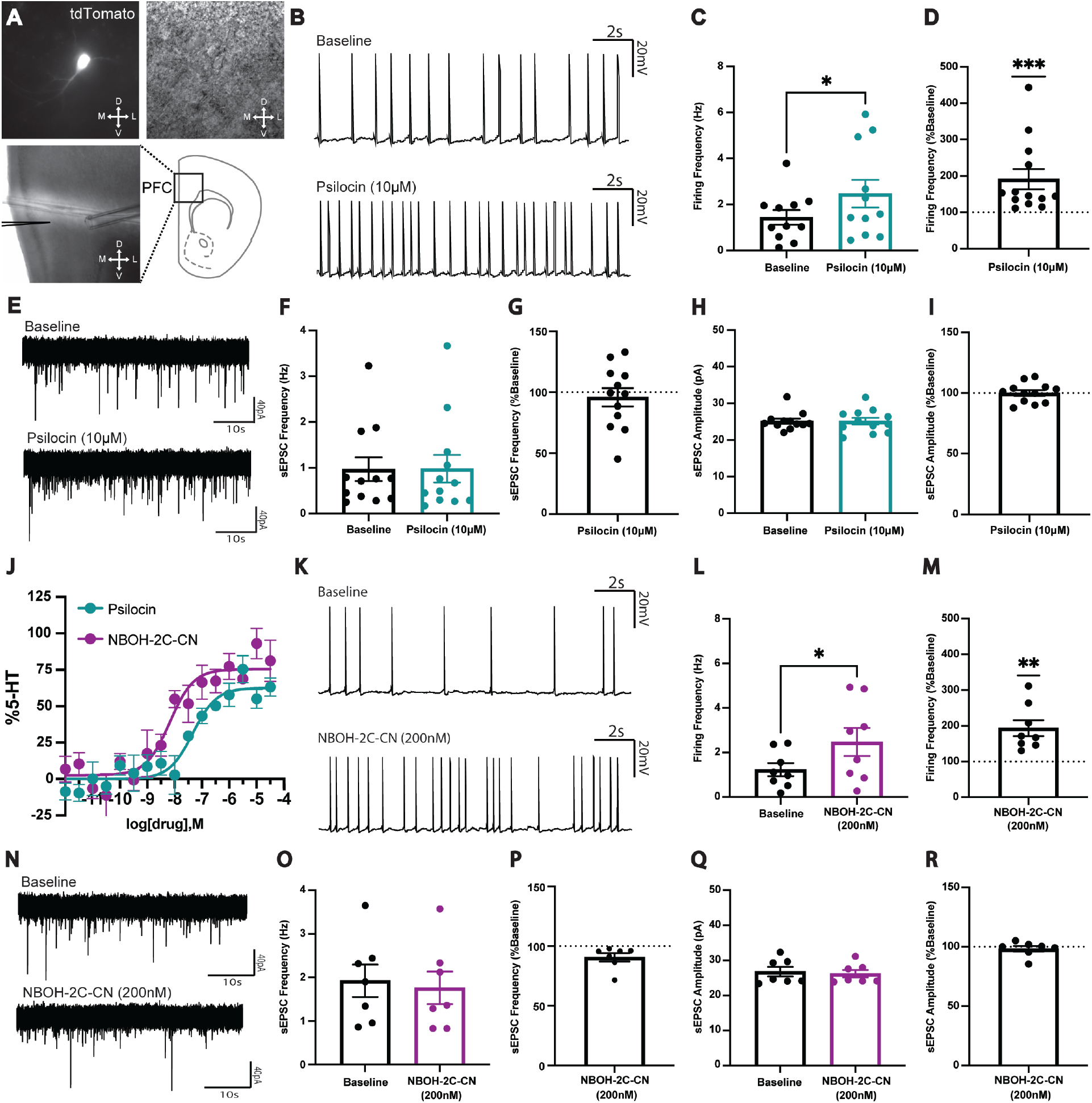
**(a)** Representative 5HT_2A_R expressing L5P neuron in PFC and its location within slice. **(b)** Representative recording of 5HT_2A_R L5P neuronal firing after psilocin (10*μ*M). **(c)** Averaged psilocin effects on firing in 5HT_2A_R L5P neurons (Data represented as mean ± SEM, n=11. Paired t-test *p<0.05). **(d)** Normalized psilocin effects on firing in 5HT_2A_R L5P neurons (Data represented as mean ± SEM, n=11. Wilcoxon signed rank test ***p (two-tailed) <0.001. **(e)** Representative recording of 5HT_2A_R L5P spontaneous excitatory post synaptic currents (sEPSCs) after psilocin (10*μ*M). **(f)** Averaged psilocin effect on sEPSC frequency (Data represented as mean ± SEM, n=12). **(g)** Normalized psilocin effect on sEPSC frequency (Data represented as mean ± SEM, n=12). **(h)** Average of psilocin effect on sEPSC amplitude (Data represented as mean ± SEM, n=12). **(i)** Normalized psilocin effect on sEPSC amplitude (Data represented as mean ± SEM, n=12). **(j)** Gq dissociation concentration-response curves at mouse 5HT_2A_R for psychedelic drugs psilocin and NBOH-2C-CN. **(k)** Representative recording of 5HT_2A_R L5P neuronal firing after NBOH-2C-CN (200nM). **(l)** Averaged NBOH-2C-CN effects on firing in 5HT_2A_R L5P neurons (Data represented as mean ± SEM, n=8. Paired t-test *p<0.05). **(m)** Normalized NBOH-2C-CN effects on firing in 5HT_2A_R L5P neurons (Data represented as mean ± SEM, n=8. One sample t-test **p (two-tailed) <0.01). **(n)** Representative recording of 5HT_2A_R L5P sEPSCs after NBOH-2C-CN (200nM). **(o)** Averaged NBOH-2C-CN effect on sEPSC frequency (Data represented as mean ± SEM, n=7). **(p)** Normalized NBOH-2C-CN effect on sEPSC frequency (Data represented as mean ± SEM, n=7). **(q)** Average of NBOH-2C-CN effect on sEPSC amplitude (Data represented as mean ± SEM, n=7). **(r)** Normalized NBOH-2C-CN effect on sEPSC amplitude (Data represented as mean ± SEM, n=7).

To confirm the specificity of this effect at 5-HT_2A_R’s, we used the preferential 5-HT_2A_R agonist NBOH-2C-CN. To guide our dose of NBOH-2C-CN, we conducted a G*α*_q_ dissociation experiment at the mouse 5HT_2A_R. The concentration response curve showed that NBOH-2C-CN demonstrated a significant increase in potency as compared to Psilocin (46nM vs 7nM, F-test **p<0.01). A 200nM dose of NBOH-2C-CN was selected to reflect this increase in potency and minimize off-target receptor activities. To examine the effects of NBOH-2C-CN on 5-HT_2A_R layer 5 pyramidal neurons in the PFC, we performed whole-cell current-clamp recordings in the PrL PFC to measure changes in firing. Focal application of NBOH-2C-CN (200nM) significantly increased firing frequency in 5-HT_2A_R expressing neurons (n=9 cells. Paired t-test *p<0.05; **Figure 3K,L**). When firing frequencies were normalized to baseline values, firing frequency increased to ∼200% (n=8. One-sample t-test. **p (two-tailed) <0.01; **Figure 3M**). To measure changes in excitatory glutamatergic synaptic transmission, we performed whole-cell voltage-clamp recordings in 5-HT_2A_R pyramidal neurons in the PrL PFC. Focal application of NBOH-2C-CN (200nM) produced no changes in spontaneous excitatory post-synaptic current (sEPSC) frequency either in raw values (n=7. **Figure 3N,O**) or normalized values (n=7. **Figure 3P**). Similarly, psilocin produced no changes sEPSC amplitude either in raw values (n=7. **Figure 3Q**) or normalized values (n=7. **Figure 3R**). To measure changes in glutamate release isolated to the pre-synaptic terminals, we performed recordings of miniature excitatory post-synaptic currents (mEPSCs). Application of NBOH-2C-CN (200nM) produced no changes in mEPSC frequency or amplitude (**Figure S1F-I**). To measure changes in inhibitory synaptic transmission, we performed whole-cell voltage-clamp recordings in 5-HT_2A_R pyramidal neurons in the PrL PFC. Focal application of NBOH-2C-CN (200nM) produced no changes in spontaneous inhibitory post-synaptic current (sIPSC) frequency or amplitude (**Figure S2F-J**).

### 3.4 Effects of Psilocin and NBOH-2C-CN on 5HT_2A_R layer 5 pyramidal neurons in the Prefrontal Cortex are not 5-HT_2C_R mediated

To specifically examine 5-HT_2C_R involvement in mediating the effects of psilocin on 5-HT_2A_R layer 5 pyramidal neurons in the PFC, we performed whole-cell current-clamp recordings in the PrL PFC to measure changes in firing. Focal application of the 5-HT_2C_R specific antagonist RS102221 (5*μ*M) significantly decreased basal firing frequency in 5-HT_2A_R expressing neurons (n=11. Paired t-test **p<0.05; **Figure 4A,B**). When firing frequencies were normalized to baseline values, firing frequency decreased to ∼30% (n=11. One-sample t-test. ****p (two-tailed) <0.0001; **Figure 4C**). Subsequent application of psilocin (10*μ*M) in the presence of the specific 5-HT_2C_R specific antagonist RS102221 (5*μ*M) significantly increased firing frequency in 5-HT_2A_R expressing neurons (n=11. Wilcoxon t-test ***p<0.001; **Figure 4A,D**). When firing frequencies were normalized to baseline values, firing frequency increased to ∼200% (n=10. One-sample t-test. *p (two-tailed) <0.05; **Figure 4E**).

**Figure 4:**
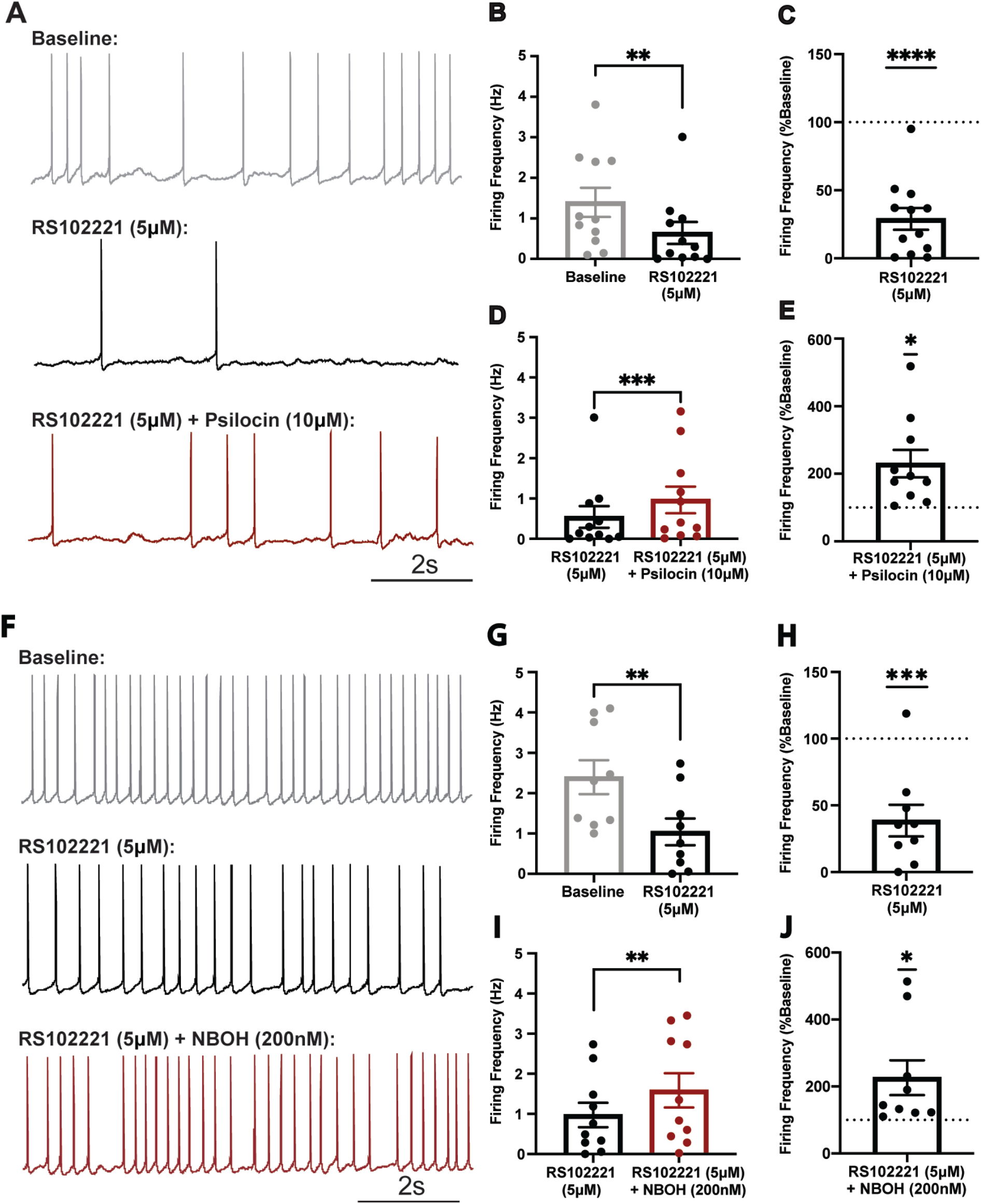
**(a)** Representative recording of 5HT_2A_R L5P neuronal firing after RS102221 (5*μ*M) and psilocin (10*μ*M). **(b)** Averaged RS102221 effects on firing in 5HT_2A_R L5P neurons (Data represented as mean ± SEM, n=11. Paired t-test **p<0.01). **(c)** Normalized RS102221 effects on firing in 5HT_2A_R L5P neurons (Data represented as mean ± SEM, n=11. One sample t-test ****p (two-tailed) <0.0001). **(d)** Averaged RS102221 + psilocin effects on firing in 5HT_2A_R L5P neurons (Data represented as mean ± SEM, n=11. Wilcoxon’s t-test ***p<0.001). **(e)** Normalized RS102221 + psilocin effects on firing in 5HT_2A_R L5P neurons (Data represented as mean ± SEM, n=10. One sample t-test *p (two-tailed) <0.05). **(f)** Representative recording of 5HT_2A_R L5P neuronal firing after RS102221 (5*μ*M) and NBOH-2C-CN (200nM). **(g)** Averaged RS102221 effects on firing in 5HT_2A_R L5P neurons (Data represented as mean ± SEM, n=9. Paired t-test **p<0.01). **(h)** Normalized RS102221 effects on firing in 5HT_2A_R L5P neurons (Data represented as mean ± SEM, n=9. One sample t-test ***p (two-tailed) <0.001). **(i)** Averaged RS102221 + NBOH-2C-CN effects on firing in 5HT_2A_R L5P neurons (Data represented as mean ± SEM, n=10. Paired t-test **p<0.01). **(j)** Normalized RS102221 + NBOH-2C-CN effects on firing in 5HT_2A_R L5P neurons (Data represented as mean ± SEM, n=9. One sample t-test *p (two-tailed) <0.05).

Similarly, to specifically examine 5-HT_2C_R involvement in mediating the effects of NBOH-2C-CN on 5-HT_2A_R layer 5 pyramidal neurons in the PFC, we performed whole-cell current-clamp recordings in the PrL PFC to measure changes in firing. Focal application of the 5-HT_2C_R specific antagonist RS102221 (5*μ*M) again significantly decreased firing frequency in 5-HT_2A_R expressing neurons (n=9. Paired t-test **p<0.01; **Figure 4F,G**). When firing frequencies were normalized to baseline values, firing frequency decreased to ∼35% (n=9. One-sample t-test. ***p (two-tailed) <0.001; **Figure 4H**). Subsequent application of NBOH-2C-CN (200nM) in the presence of the specific 5-HT_2C_R specific antagonist RS102221 (5*μ*M) significantly increased firing frequency in 5-HT_2A_R expressing neurons (n=10. Paired t-test **p<0.01; **Figure 4F,I**). When firing frequencies were normalized to baseline values, firing frequency increased to ∼200% (n=9. One-sample t-test. *p (two-tailed) <0.05; **Figure 4J**).

### 3.5 Effects of Psilocin and NBOH-2C-CN on 5HT_2A_R layer 5 pyramidal neurons in the Prefrontal Cortex are 5-HT_2A_R mediated

To specifically examine 5-HT_2A_R involvement in mediating the effects of psilocin on 5-HT_2A_R layer 5 pyramidal neurons in the PFC, we performed whole-cell current-clamp recordings in the PrL PFC to measure changes in firing. Focal application of the 5-HT_2A_R specific antagonist M100907 (200nM) significantly decreased basal firing frequency in 5-HT_2A_R expressing neurons (n=8. Paired t-test *p<0.05; **Figure 5A,B**). When firing frequencies were normalized to baseline values, firing frequency decreased to ∼40% (n=8. One-sample t-test. **p=0.0071; **Figure 5C**). Subsequent application of psilocin (10*μ*M) in the presence of the specific 5-HT_2A_R specific antagonist M100907 (200nM) had no effect on firing frequency in 5-HT_2A_R expressing neurons (n=10. **Figure 5A,D**). When firing frequencies were normalized to baseline values, again no effect on firing frequency was observed (n=10. **Figure 5E**).

**Figure 5:**
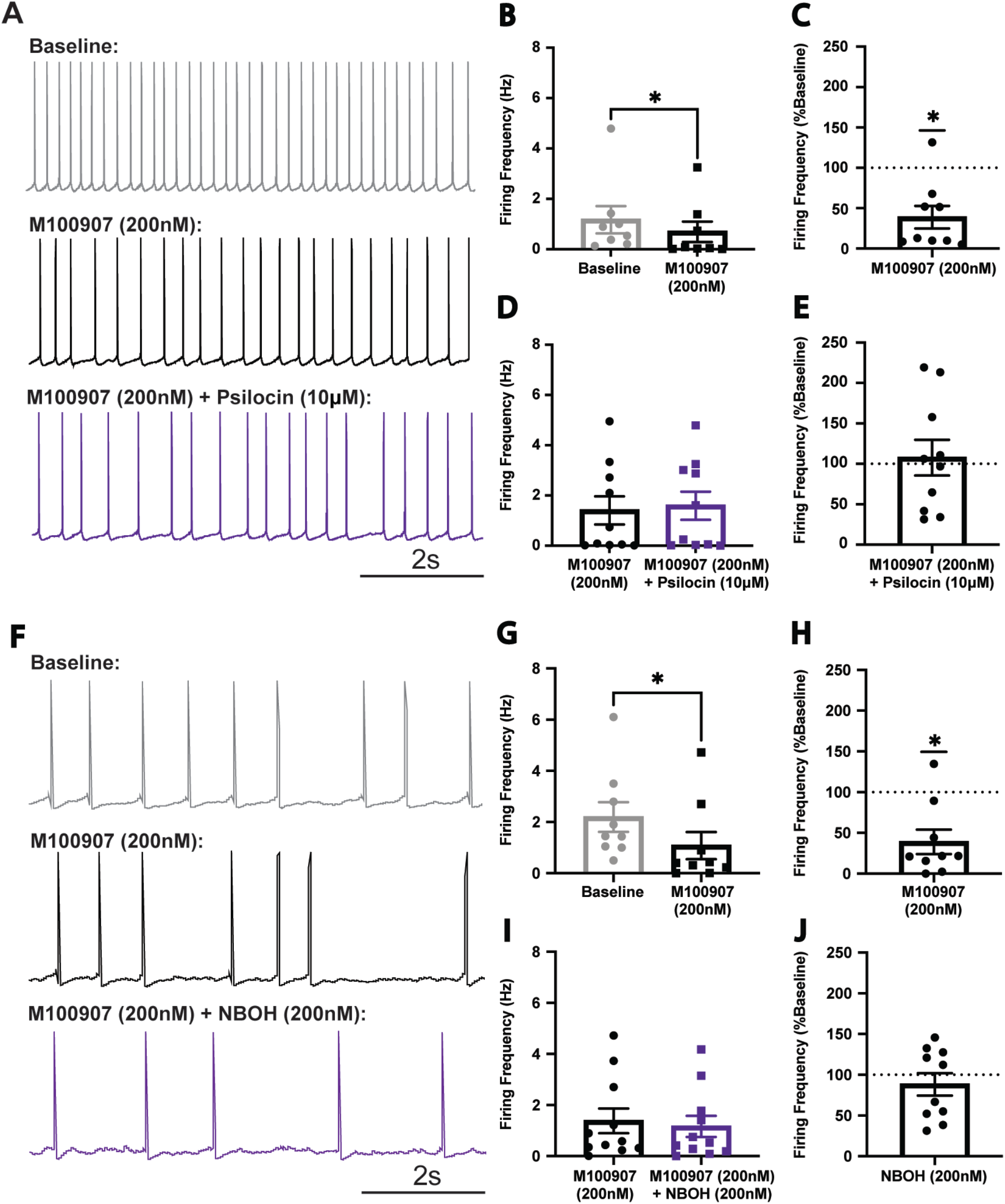
**(a)** Representative recording of 5HT_2A_R L5P neuronal firing after M100907 (200nM) and psilocin (10*μ*M). **(b)** Averaged M100907 effects on firing in 5HT_2A_R L5P neurons (Data represented as mean ± SEM, n=8. Paired t-test *p<0.05). **(c)** Normalized M100907 effects on firing in 5HT_2A_R L5P neurons (Data represented as mean ± SEM, n=8. Wilcoxon signed rank t-test *p (two-tailed) <0.05). **(d)** Averaged M100907 + psilocin effects on firing in 5HT_2A_R L5P neurons (Data represented as mean ± SEM, n=10). **(e)** Normalized M100907 + psilocin effects on firing in 5HT_2A_R L5P neurons (Data represented as mean ± SEM, n=10). **(f)** Representative recording of 5HT_2A_R L5P neuronal firing after M100907 (200nM) and NBOH-2C-CN (200nM). **(g)** Averaged M100907 effects on firing in 5HT_2A_R L5P neurons (Data represented as mean ± SEM, n=9. Wilcoxon t-test *p<0.05). **(h)** Normalized M100907 effects on firing in 5HT_2A_R L5P neurons (Data represented as mean ± SEM, n=9. Wilcoxon signed rank t-test *p (two-tailed) <0.01). **(i)** Averaged M100907 + NBOH-2C-CN effects on firing in 5HT_2A_R L5P neurons (Data represented as mean ± SEM, n=11). **(j)** Normalized M100907 + NBOH-2C-CN effects on firing in 5HT_2A_R L5P neurons (Data represented as mean ± SEM, n=10).

Similarly, to examine 5-HT_2A_R involvement in mediating the effects of NBOH-2C-CN on 5-HT_2A_R layer 5 pyramidal neurons in the PFC, we performed whole-cell current-clamp recordings in the PrL PFC to measure changes in firing. Focal application of the 5-HT_2A_R specific antagonist M100907 (200nM) again significantly decreased firing frequency in 5-HT_2A_R expressing neurons (n=9. Wilcoxon t-test *p<0.05; **Figure 5F,G**). When firing frequencies were normalized to baseline values, firing frequency decreased to ∼35% (n=9. Wilcoxon signed rank t-test. *p (two-tailed) <0.05; **Figure 5H**). Subsequent application of NBOH-2C-CN (200nM) in the presence of the specific 5-HT_2A_R specific antagonist M100907 (200nM) had no effect on firing frequency in 5-HT_2A_R expressing neurons (n=11. **Figure 5F,I**). When firing frequencies were normalized to baseline values, again no effect on firing frequency was observed (n=10. **Figure 5J**).

### 3.6 Effects of Psilocin and NBOH-2C-CN on 5HT_2A_R layer 5 pyramidal neurons in the Prefrontal Cortex are G*α*_q_ mediated

Given that 5-HT_2A_ receptors are G protein-coupled receptors (GPCRs) known to signal through the G*α*_q_ family of heterotrimeric G proteins and recruit β-arrestin (βArr) proteins, we performed whole-cell current-clamp recordings in the PrL PFC to measure changes in firing induced by psilocin and NBOH-2C-CN in the presence of the selective G*α*_q_ inhibitor FR900359. Focal application of the G*α*_q_ inhibitor FR900359 (1*μ*M) significantly decreased firing frequency in 5-HT_2A_R expressing neurons (n=10. Paired t-test **p<0.01; **Figure 6A,B**). When firing frequencies were normalized to baseline values, firing frequency decreased to ∼40% (n=10. Wilcoxon signed rank t-test. **p<0.01; **Figure 6C**). Subsequent application of psilocin (10*μ*M) in the presence of the selective G*α*_q_ inhibitor FR900359 (1*μ*M) did not significantly change firing frequency in 5-HT_2A_R expressing neurons (n=10. **Figure 6A,D**). When firing frequencies were normalized to baseline values, again no effect on firing frequency was observed (n=10. **Figure 6E**).

**Figure 6:**
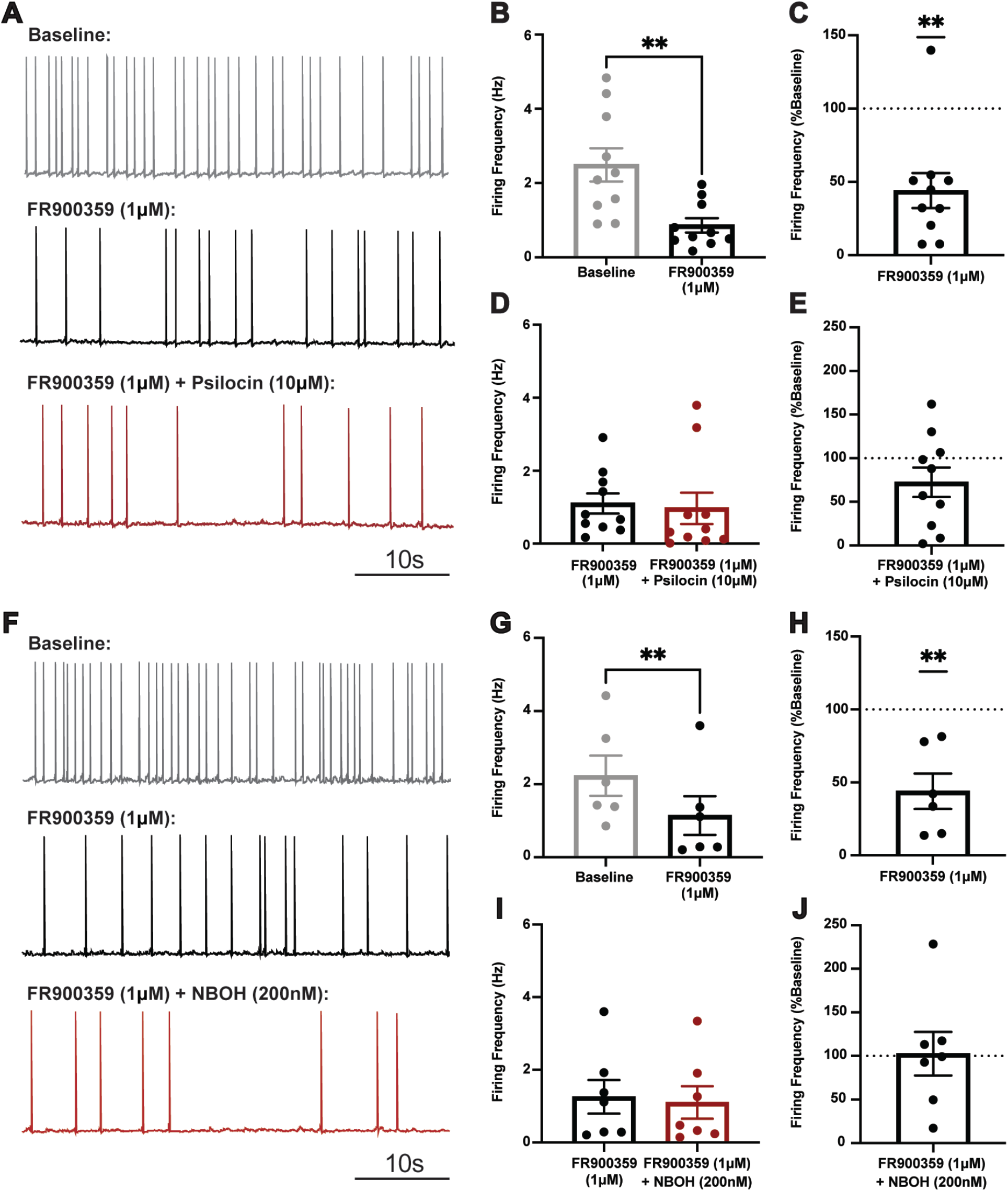
**(a)** Representative recording of 5HT_2A_R L5P neuronal firing after FR900359 (1*μ*M) and psilocin (10*μ*M) showing significant change in frequency. **(b)** Averaged FR900359 effects on firing in 5HT_2A_R L5P neurons (Data represented as mean ± SEM, n=10. Paired t-test **p<0.01). **(c)** Normalized FR900359 effects on firing in 5HT_2A_R L5P neurons (Data represented as mean ± SEM, n=10. Wilcoxon signed rank t-test **p (two-tailed) <0.01). **(d)** Averaged FR900359 + psilocin effects on firing in 5HT_2A_R L5P neurons (Data represented as mean ± SEM, n=10). **(e)** Normalized FR900359 + psilocin effects on firing in 5HT_2A_R L5P neurons (Data represented as mean ± SEM, n=10). **(f)** Representative recording of 5HT_2A_R L5P neuronal firing after FR900359 (1*μ*M) and NBOH-2C-CN (200nM) showing significant change in frequency. **(g)** Averaged FR900359 effects on firing in 5HT_2A_R L5P neurons (Data represented as mean ± SEM, n=6. Paired t-test **p<0.01). **(h)** Normalized FR900359 effects on firing in 5HT_2A_R L5P neurons (Data represented as mean ± SEM, n=6. One sample t-test **p (two-tailed) <0.01). **(i)** Averaged FR900359 + NBOH-2C-CN effects on firing in 5HT_2A_R L5P neurons (Data represented as mean ± SEM, n=7). **(j)** Normalized FR900359 + NBOH-2C-CN effects on firing in 5HT_2A_R L5P neurons (Data represented as mean ± SEM, n=7).

Similarly, to examine G*α*_q_ signaling in mediating the effects of NBOH-2C-CN on 5-HT_2A_R layer 5 pyramidal neurons in the PFC, we performed whole-cell current-clamp recordings in the PrL PFC to measure changes in firing. Focal application of the G*α*_q_ inhibitor FR900359 (1*μ*M) again decreased firing frequency in 5-HT_2A_R expressing neurons (n=6. Paired t-test **p<0.01; **Figure 6F,G**). When firing frequencies were normalized to baseline values, firing frequency decreased to ∼40% (n=6. One-sample t-test. **p<0.01; **Figure 6H**). Subsequent application of NBOH-2C-CN (200nM) in the presence of the selective G*α*_q_ inhibitor FR900359 (1*μ*M) did not significantly change firing frequency in 5-HT_2A_R expressing neurons (n=7. Paired t-test *p<0.05; **Figure 6F,I**). When firing frequencies were normalized to baseline values, again no effect on firing frequency was observed (n=7. One-sample t-test. **p (two-tailed) <0.01; **Figure 6J**).

### 3.7 Intracellular application of a G*α*_q_ inhibitor blocks the effects of psilocin and NBOH-2C-CN on 5HT_2A_R layer 5 pyramidal neurons in the Prefrontal Cortex

To further validate that psilocin exerts its effect on 5HT_2A_R layer 5 pyramidal neurons in the Prefrontal Cortex via a G*α*_q_ signaling mechanism, we included the selective G*α*_q_ inhibitor FR900359 (1*μ*M) in the internal solution of the recording pipette and performed whole-cell current-clamp recordings in the PrL PFC to measure changes in firing induced by psilocin and NBOH-2C-CN. Focal application of psilocin (10*μ*M) significantly and paradoxically reduced firing frequency in 5-HT_2A_R expressing neurons (n=9. Paired t-test *p<0.05; **Figure 7A,B**). When firing frequencies were normalized to baseline values, firing frequency was significantly reduced (n=10. One-sample t-test. *p (two-tailed) <0.05; **Figure 7C**).

**Figure 7:**
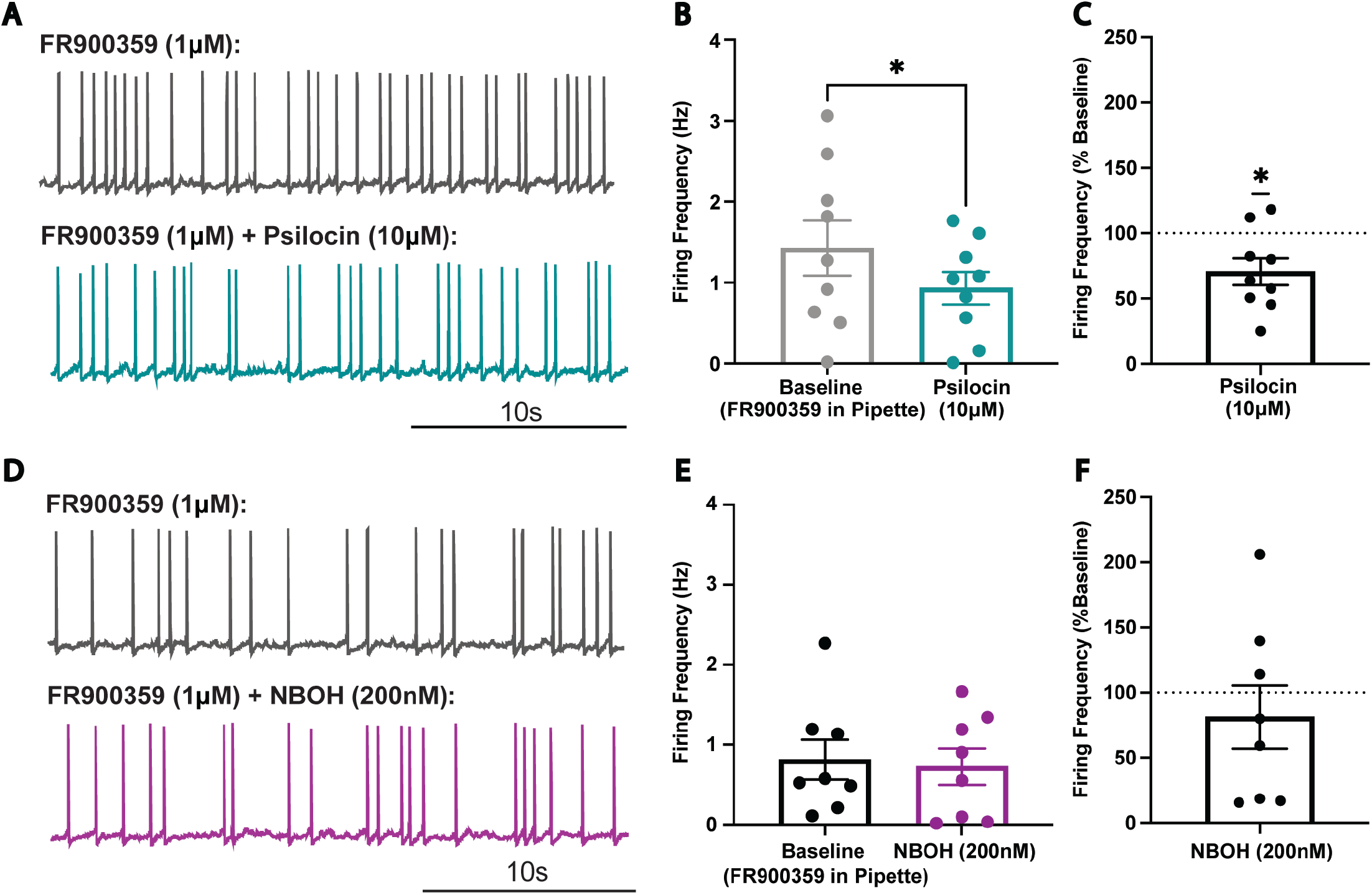
**(a)** Representative recording of 5HT_2A_R L5P neuronal firing with FR900359 (1*μ*M) in internal solution and psilocin (10*μ*M) showing significant change in frequency. **(b)** Averaged FR900359 + psilocin effects on firing in 5HT_2A_R L5P neurons (Data represented as mean ± SEM, n=9. Paired t-test **p<0.05). **(c)** Normalized FR900359 + psilocin effects on firing in 5HT_2A_R L5P neurons (Data represented as mean ± SEM, n=10. One sample t-test *p (two-tailed) <0.05). **(d)** Representative recording of 5HT_2A_R L5P neuronal firing with FR900359 (1*μ*M) in internal solution and NBOH-2C-CN (200nM). **(e)** Averaged FR900359 + NBOH-2C-CN effects on firing in 5HT_2A_R L5P neurons (Data represented as mean ± SEM, n=8.). **(f)** Normalized FR900359 + NBOH-2C-CN effects on firing in 5HT_2A_R L5P neurons (Data represented as mean ± SEM, n=8).

Focal application of NBOH-2C-CN (200nM) did not significantly change firing frequency in 5-HT_2A_R expressing neurons exposed to FR900359 (1*μ*M) (n=8. **Figure 7D,E**). When firing frequencies were normalized to baseline values, no change was observed (n=8. **Figure 7F**).

## 4. Discussion

The major finding of this paper is that the firing frequency of layer 5 pyramidal neurons expressing the 5-HT_2A_R identified using a transgenic mouse line targeting 5-HT_2A_R neurons via a tamoxifen-sensitive tdTomato reporter is robustly and consistently increased by the psychedelic compound psilocin. These findings are in contrast to the variable effects observed in unspecified layer 5 pyramidal neurons. The increase in firing in 5-HT_2A_R neurons was also observed with application of the 5-HT_2A_R selective agonist 25-CN-NBOH. Importantly, neither psilocin nor 25-CN-NBOH was found to significantly affect excitatory or inhibitory synaptic transmission, suggesting that the excitatory effects on 5-HT_2A_R layer 5 pyramidal neurons are due to postsynaptic actions and not changes in synaptic input. The excitatory effects were found to be mediated by 5-HT_2A_Rs as blockade of the 5-HT_2A_R prevented the increase in firing frequency caused by psilocin or 25-CN-NBOH. In contrast, we found no contribution of 5-HT_2C_Rs as blockade of the 5-HT_2C_R did not prevent the increase in firing frequency caused by psilocin or 25-CN-NBOH. Interestingly, both 5-HT_2A_Rs and 5-HT_2C_Rs mediate tonic firing of 5-HT_2A_R PFC pyramidal neurons as reductions in firing frequency were observed with both 5-HT_2A_R and 5-HT_2C_R antagonists. Furthermore, we show that the increase in firing activity by psilocin is mediated by an intracellular G*α*_q_ signaling pathway downstream of the 5-HT_2A_R, as application of a selective G*α*_q_ inhibitor blocked the increase in firing frequency caused by psilocin or 25-CN-NBOH. Given the highly interconnected nature of cortical pyramidal neurons, we propose that this G*α*_q_ mediated activation of 5-HT_2A_Rs increases firing in 5-HT_2A_R layer 5 pyramidal neurons which in turn promotes glutamatergic signaling onto other layer 5 pyramidal neurons as well as output regions. Collectively, these studies identify important population-specific neurobiological mechanisms underlying the effects of psilocin on the prefrontal cortex.

Previous studies have shown that in layer 5 pyramidal neurons, the most robust cellular effect of 5-HT_2A_R activation is an increase in the frequency and amplitude of glutamatergic signaling^40, 41^. However, 5-HT_2A_R activation was also reported not to excite pyramidal neurons leading some investigators to propose that postsynaptic 5-HT_2A_Rs on pyramidal neurons trigger the release of glutamate from thalamocortical fibers via an unidentified retrograde messenger. In the present study, we found that psilocin exerted differential effects on firing in layer 5 pyramidal neurons in C57/B6J mice. These results are consistent with prior studies showing increased firing in only a subset of layer 5 pyramidal neurons both *in vivo* and *ex vivo* consistent with the idea that psychedelics exert differential effects across cortical regions depending on specific drug, dose of drug, and 5HT_2A_R density in those neuronal populations^42-44^. Importantly, these prior studies examined pyramidal neuron activity in an unspecified manner, and no study directly targeted the 5-HT_2A_R pyramidal cell population. In the present study we employed a transgenic reporter mouse to specifically target 5-HT_2A_R pyramidal neurons and found that psilocin and the 5-HT_2A_R-selective agonist 25-CN-NBOH significantly increased the firing frequency of 5-HT_2A_R pyramidal neurons in the PFC suggesting that the 5-HT_2A_R population represents a PFC cell population that is uniquely sensitive to the actions of psychedelics and may be the source of increased glutamate release broadly in the PFC.

Interestingly, we found that neither psilocin nor 25-CN-NBOH had a significant effect on the excitatory or inhibitory synaptic transmission of 5-HT_2A_R pyramidal neurons. This finding suggests that the increase in firing frequency of 5-HT_2A_R pyramidal neurons in the PFC results from a direct effect of psilocin on these neurons as opposed to changes in intrinsic or extrinsic excitatory or inhibitory transmission. Our results therefore are inconsistent with the involvement of a retrograde tracer and instead suggest that psilocin and other psychedelics excite 5-HT_2A_R layer 5 pyramidal neurons causing an increase in glutamatergic recurrent network activity. Indeed, one recent study showed that inhibition of post-synaptic 5HT_2A_Rs in transfected neurons failed to inhibit increases in sEPSCs^45^, a result also inconsistent with the idea that postsynaptic 5HT_2A_Rs signal the release of a retrograde messenger capable of inducing glutamate release. In the present study, we provide evidence for the simpler explanation that 5-HT_2A_R layer 5 pyramidal neurons in the PFC are a discrete subpopulation capable of increasing the sEPSC activity of nearby non-5-HT_2A_R layer 5 pyramidal neurons.

5-HT_2A_R agonism has been shown to be crucial in mediating the psychoactive effects of psychedelics in preclinical models, and recent human imaging studies have elaborated upon these findings, identifying the PFC as a key brain region involved in mediating the effects of psychedelics. Human positron emission tomography (PET) studies have shown that psilocin’s psychedelic effects are specifically correlated with PFC 5-HT_2A_R occupancy^31^. Additionally, an fMRI study that found that the degree of increase in PFC activity one day post psilocin administration was predictive of treatment response measured at five weeks post treatment^46^. However, recent animal studies have disputed the involvement of the 5-HT_2A_R in mediating the antidepressant activity of therapeutically promising psychedelics showing that modest doses of ketanserin, a nonselective 5-HT_2_R antagonist, failed to block psilocybin-induced changes in hippocampal synaptic response^47^. In the present study, we demonstrate that the 5HT_2A_R is required for the increased firing activity seen in layer 5 5-HT_2A_R pyramidal neurons in the PFC as prior application of the 5-HT_2A_R antagonist M100907 blocked the effects of either psilocin or 25-CN-NBOH on firing. This suggests that effects of psilocin across different brain regions may be modulated by different receptors.

Currently available psychedelic compounds including psilocin, 25-CN-NBOH, LSD, and mescaline activate several biogenic amine receptors present in the prefrontal cortex in addition to the 5-HT_2A_R. One such target is the 5-HT_2C_R, with one study showing that knockout of the 5-HT_2C_R in mice resulted in attenuation of the head twitch response, a common measure of hallucinogenic effects in mice, in response to DOI^48^. In the current study, we show that prior application of the 5-HT_2C_R antagonist RS102221 did not prevent the increase in firing in layer 5 5-HT_2A_R pyramidal neurons seen with application of psilocin or 25-CN-NBOH suggesting that the 5-HT_2C_R does not play a role in mediating the effects of either compound. Interestingly, we note a significant reduction in firing in 5-HT_2A_R layer 5 pyramidal neurons with application of the 5-HT_2A_R antagonist M100907, 5-HT_2C_R antagonist RS102221, or the G*α*_q_ inhibitor FR900359 suggesting a role for the 5-HT_2A_R, the 5-HT_2C_R, and intracellular G*α*_q_ mediated signaling in the maintenance of tonic excitation of these neurons; however, only 5-HT_2A_R and intracellular G*α*_q_ signaling mediate the direct effects of psilocin’s excitation of 5-HT_2A_R layer 5 pyramidal neurons.

One concept that has yet to be elucidated in relation to the specific actions of psychedelic drugs is biased signaling. Biased signaling or functional selectivity is a ligand-dependent phenomenon where certain ligands are known to preferentially activate one signaling pathway. Canonically, GPCRs signal both through the direct actions of G protein mediated signaling cascades as well as through βArr-mediated signaling pathways. It has been proposed that ligands with biased profiles may be useful for activating desired (i.e. therapeutically efficacious) pathways rather than undesired (i.e. unwanted side effect producing) pathways downstream of a given GPCR^49^. Whether the hallucinogenic and therapeutic effects of serotonergic psychedelics are dissociable is an area of growing interest in the study of psychedelics, generating many novel compounds in recent years^25-27^. LSD is shown to preferentially signal through βArr2 at the 5-HT_2A_ receptor^11, 50^ and numerous LSD-elicited responses are either significantly attenuated or completely absent in βArr2-KO mice^51^. In the current study, we show that the 5-HT_2A_R mediated effects specifically in 5-HT_2A_R prefrontal cortex pyramidal neurons are dependent upon a G*α*_q_ signaling pathway. Specifically, inhibition of G*α*_q_ signaling in the slice or restricted to the recorded neuron prevented the increase in firing seen with application of psilocin or 25-CN-NBOH. Although there have been reports that psychedelic compounds differentially couple 5-HT_2A_Rs to Gα_i/o_ signaling in brain^14, 52^, our results are consistent with prior studies showing that 5-HT_2A_Rs mainly couple to pertussis-toxin insensitive Gα_q_-like transducers^50, 53-56^, and prior *in vivo* reports showing that genetic deletion of one copy of Gα_q_ abolished psychedelic drug-induced c-Fos expression^53^.

Understanding the alterations in prefrontal cortex neural activity is important in the context of psilocin’s emerging therapeutic utility. Neuropsychiatric disorders including anxiety, depression, and substance use are associated with dysregulated activity in the frontal cortex^35^ followed by eventual reductions in frontal cortex synapses^57^. Increasing the level of activity amongst serotonergic neurons may counteract such deficits, providing a mechanism for ameliorating depressive phenotypes and opposing the loss of neural connectivity in these areas seen with advanced depressive phenotypes. An important preclinical study identified that a single dose of psilocybin was shown to evoke dendritic spine growth in the frontal cortex of mice 24 hours after administration^58^. The structural changes appeared quickly and were long-lasting, persisting up to 28 days^58^. This study provides important information about the structural changes in the PFC occurring 24 hours after systemic exposure. The acute increase in activity we report here specific to the 5-HT_2A_R layer 5 pyramidal neurons provides one mechanism from which these structural changes might originate.

Preclinical studies provide the opportunity for a more detailed mechanistic assessment of psychedelic drug function without the human confounds of anticipation or expectation bias; however, there are important differences between rodent models of exposure and the human experience. While the affective components of the psychedelic experience are more difficult to mimic in rodent models, one benefit of preclinical models like the one used here is that we can more mechanistically assess the direct functional consequences of psychedelic drugs. In the present study, we chose to probe the effects of classical hallucinogens as this study focused on the 5-HT_2A_R mediated component of psychedelic function. Given that receptor pharmacologies vary among different psychedelic drugs, future studies should examine the effects of other hallucinogenic psychedelics as well as non-hallucinogenic 5-HT_2A_R agonists. The effects of psychedelic drugs have been shown to be altered in naturally occurring single nucleotide polymorphisms in the 5-HT2A receptor gene^59^. Of note, although the mouse 5-HT_2A_R and the human 5-HT_2A_R sequences are highly conserved, there are possible functionally relevant differences^60^ that could impact drug effects and the potential effects of these sequence differences should also be examined. In addition, it will be important to identify specific circuit changes involved in psilocin’s action including those involving the PFC. Mapping the distinct projections from 5-HT_2A_R layer 5 pyramidal neurons in the PFC using the 5-HT_2A_R-reporter mouse and viral tracing approaches will also be important as it will allow us to identify circuit-specific sequelae of psilocin-induced changes in 5-HT_2A_R pyramidal neuron activity. Our work suggests that psychedelic drug-induced changes in activity patterns are complex and may involve differential network effects, and future work should also measure changes in other 5-HT_2A_R expressing neuron populations. Additionally, it will be important to investigate the impact of other neuromodulatory systems and co-expression studies will help us to better understand distinct subpopulations of 5-HT_2A_R pyramidal neurons. One study found that *μ*-opioid receptor activation completely blocked the ability of 5-HT_2A_R agonists to excite any layer 5 pyramidal neurons^61^, presumably including those that express the 5-HT_2A_R like the ones in the present study, suggesting that other systems may be involved in psychedelic drug effects.

There has recently been a renewed interest in the scientific understanding and translational potential of psilocin and other psychedelic drugs. Essential for the continued development of clinical applications however is a clear, detailed understanding of how psychedelics exert their effects. In the current study, we detail the effects of psilocin in a key brain region highly relevant to human disease and identify population-specific effects that are mediated by a G*α*_q_ signaling pathway downstream of the 5-HT_2A_R. Additionally, we identify 5-HT_2A_R layer 5 pyramidal neurons in the PFC as a discrete subpopulation capable of increasing the sEPSC activity of nearby non-5-HT_2A_R layer 5 pyramidal neurons.

---

**Figure S1:**
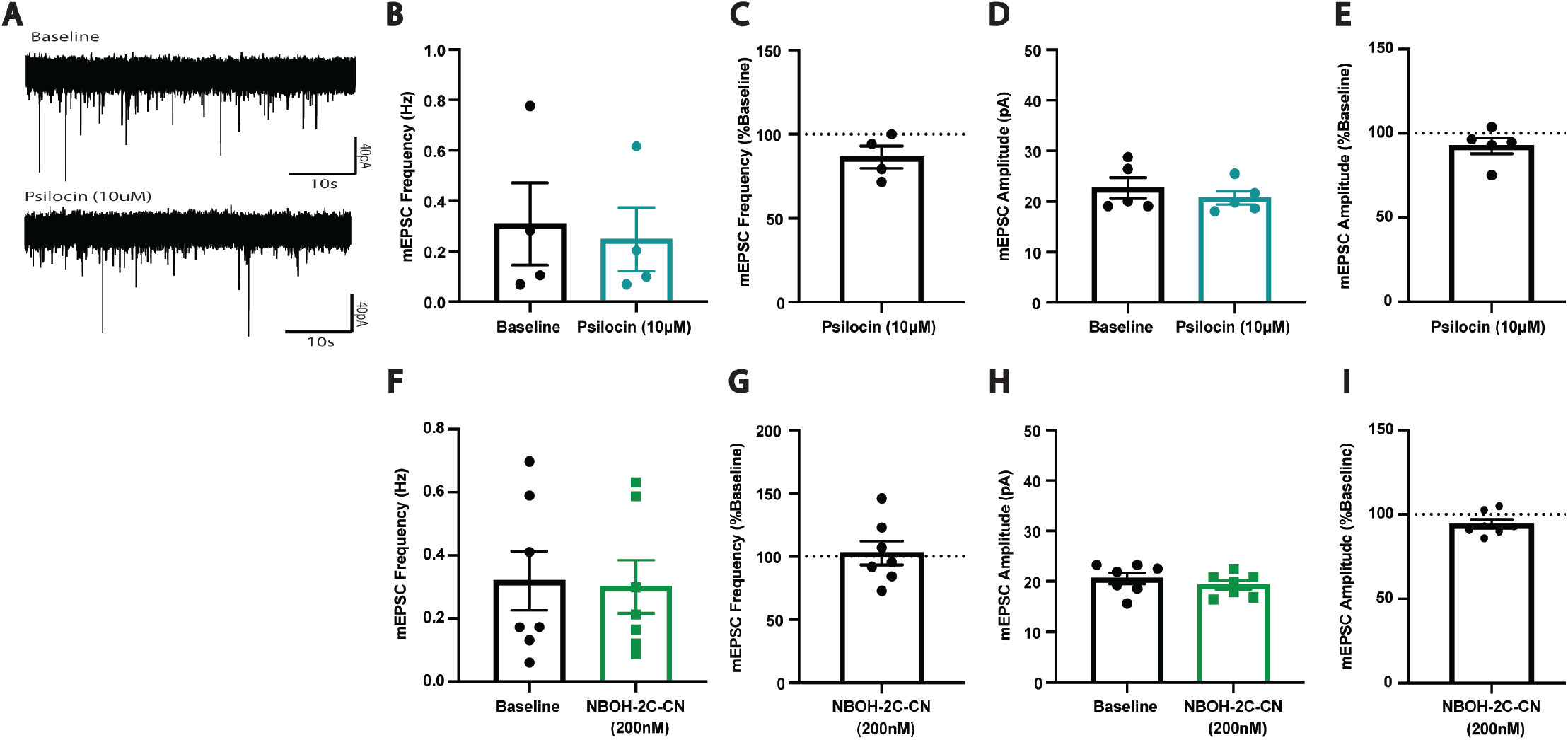
**(a)** Representative recording of 5HT_2A_R L5P miniature excitatory post synaptic currents (mEPSCs) after psilocin (10*μ*M). **(b)** Averaged psilocin effect on mEPSC frequency (Data represented as mean ± SEM, n=4). **(c)** Normalized psilocin effect on mEPSC frequency (Data represented as mean ± SEM, n=4). **(d)** Average of psilocin effect on mEPSC amplitude (Data represented as mean ± SEM, n=4). **(e)** Normalized psilocin effect on mEPSC amplitude (Data represented as mean ± SEM, n=4). **(f)** Averaged NBOH-2C-CN effect on mEPSC frequency (Data represented as mean ± SEM, n=7). **(g)** Normalized NBOH-2C-CN effect on mEPSC frequency (Data represented as mean ± SEM, n=7). **(h)** Average of NBOH-2C-CN effect on mEPSC amplitude (Data represented as mean ± SEM, n=7). **(i)** Normalized NBOH-2C-CN effect on mEPSC amplitude (Data represented as mean ± SEM, n=7).

**Figure S2:**
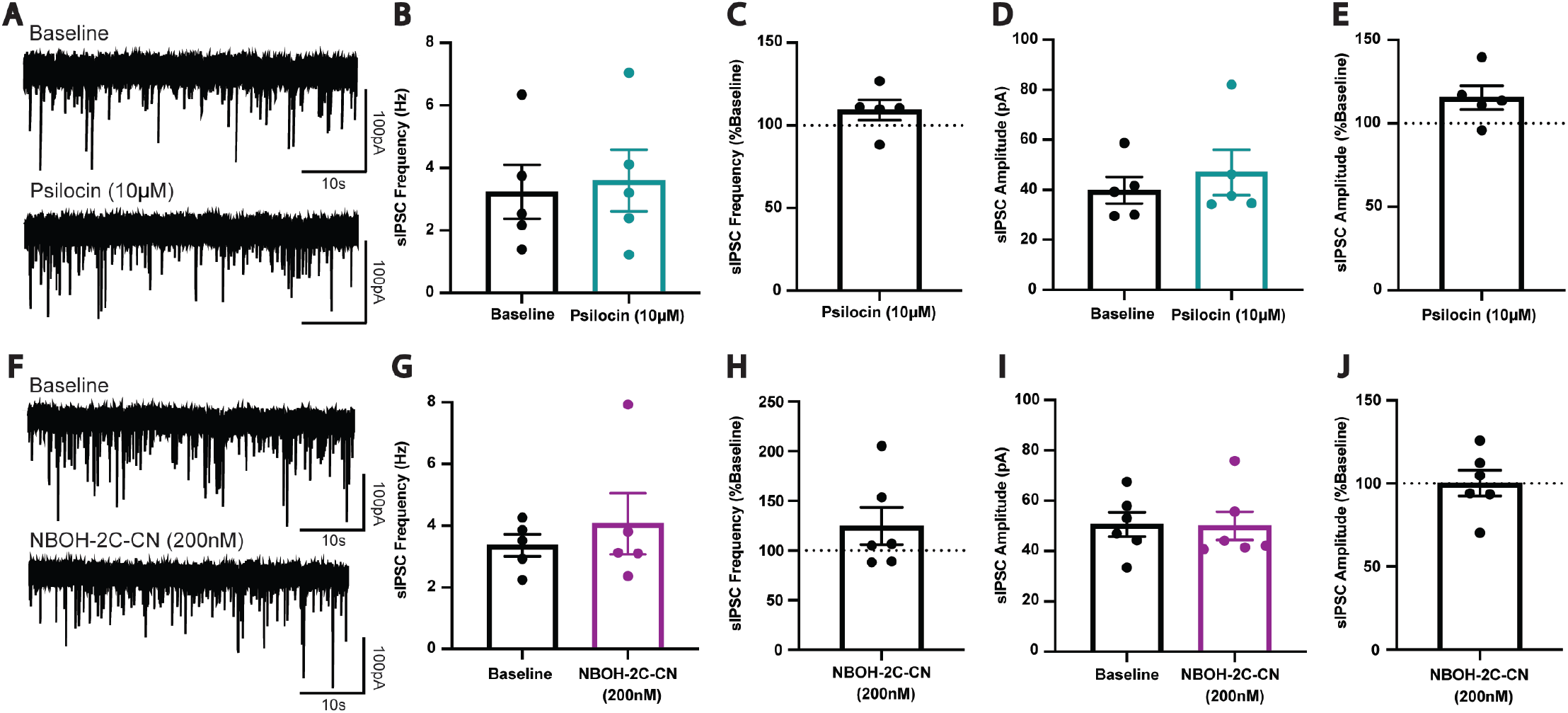
**(a)** Representative recording of 5HT_2A_R L5P spontaneous inhibitory post synaptic currents (sIPSCs) after psilocin (10*μ*M). **(b)** Averaged psilocin effect on sIPSC frequency (Data represented as mean ± SEM, n=5). **(c)** Normalized psilocin effect on sIPSC frequency (Data represented as mean ± SEM, n=5). **(d)** Average of psilocin effect on sIPSC amplitude (Data represented as mean ± SEM, n=5). **(e)** Normalized psilocin effect on sIPSC amplitude (Data represented as mean ± SEM, n=5). **(f)** Representative recording of 5HT_2A_R L5P spontaneous inhibitory post synaptic currents (sIPSCs) after NBOH-2C-CN (200nM). **(g)** Averaged NBOH-2C-CN effect on sIPSC frequency (Data represented as mean ± SEM, n=5). **(h)** Normalized NBOH-2C-CN effect on sIPSC frequency (Data represented as mean ± SEM, n=5). **(i)** Average of NBOH-2C-CN effect on sIPSC amplitude (Data represented as mean ± SEM, n=5). **(j)** Normalized NBOH-2C-CN effect on sIPSC amplitude (Data represented as mean ± SEM, n=5).

